# Progressive hypothalamic neuroinflammation after ovariectomy in mice parallels age-related transcriptomic changes in the female human hypothalamus

**DOI:** 10.1101/2024.10.05.616758

**Authors:** Jordana C. B. Bloom, Encarnación Torres, Sidney A. Pereira, Liliana Arvizu-Sanchez, Audrey N. Fontes, Hadine Joffe, David C. Page, Victor M. Navarro

**Author notes:** These authors contributed equally to this work (Encarnación Torres, Sidney A. Pereira).

## Abstract

The hypothalamic changes that occur after the loss of ovarian estrogen remain poorly characterized. Here, we performed a comprehensive temporal characterization of the mouse hypothalamus following ovariectomy (OVX), combining physiological measurements with bulk RNA-sequencing of the posterior hypothalamus (PH) and preoptic area (POA) at short-term (14 days) and long-term (4 months) post-OVX. Serum LH levels rose progressively and then declined, while core temperature peaked early and subsequently normalized, recapitulating the endocrine and thermoregulatory dynamics of reproductive aging in humans. Transcriptomic analysis revealed time-dependent activation of inflammatory pathways, glial markers, and KNDy neuron-related gene networks, with the most pronounced changes emerging at 4 months post-OVX, particularly in the PH. Immunofluorescence confirmed increased NKB release, declining KNDy neuronal activity, and heightened astrocytic reactivity in the arcuate nucleus after prolonged estrogen withdrawal. To contextualize these findings, we analyzed publicly available human hypothalamic RNA-seq data across chronological age. Age-related transcriptomic patterns in women, including progressive inflammatory signaling, glial activation, and altered KNDy gene expression, showed significant correlation with the OVX mouse model, particularly at the pathway level. These findings establish a temporal framework for hypothalamic molecular changes after estrogen withdrawal, identify conserved neuroinflammatory signatures across species, and provide a preclinical platform for testing interventions targeting menopausal-associated hypothalamic dysfunction.

## Introduction

Menopause represents a significant life transition for women, characterized by physiological changes extending beyond reproductive function^1^. Understanding the neurobiological basis of these changes is crucial for advancing women’s health and developing targeted interventions for menopausal symptoms^2^. While much attention has been focused on ovarian function, research focused on characterizing the changes that happen at the level of the brain, particularly the hypothalamus, has lagged. The hypothalamus contains neuroendocrine centers that both control and respond to estrogens^3,4^, and hypothalamic changes are implicated in disrupted thermoregulation, sleep patterns, and increased mood disorders during menopause^5–12^.

Hypothalamic KNDy neurons (expressing kisspeptin, neurokinin B, and dynorphin A) have emerged as central players in vasomotor symptoms (VMS), including hot flashes and night sweats, which are among the most common and disruptive menopausal symptoms^13,14^. The identification of KNDy neurons as VMS triggers has opened new avenues for understanding and treating these symptoms^13–15^, ultimately leading to the FDA approval of NK3R antagonists for VMS^16^. However, our knowledge of broader hypothalamic changes during and after estrogen withdrawal remains incomplete, compounded by limitations in current animal models^17^.

A critical gap in the field has been the absence of a well-characterized temporal model of hypothalamic changes following estrogen loss. Most ovariectomy (OVX) studies in rodents examine a single short-term timepoint, typically 1–3 weeks post-surgery, which captures the acute response to estrogen withdrawal but fails to model the progressive changes that develop over months to years in women. This is particularly relevant because VMS and other menopausal symptoms show distinct temporal trajectories. VMS typically peak during the perimenopausal period and gradually resolve years after menopause, while inflammatory and neurodegenerative processes may continue to progress^18^. Whether these clinically distinct phases correspond to discrete molecular programs in the hypothalamus has not been systematically investigated.

Here, we address this gap by performing a comprehensive temporal analysis of the mouse hypothalamus at multiple timepoints after OVX, integrating endocrine measurements, core body temperature, bulk RNA-sequencing of two hypothalamic subregions, and protein-level validation. We then leverage publicly available human hypothalamic transcriptomic data from the GTEx consortium to assess whether the mouse molecular signatures parallel age-related changes observed in women across the adult lifespan, while acknowledging the inherent limitations of the human observational dataset. By combining rigorous experimental data with complementary human observations, this study provides both mechanistic insight and a preclinical platform for developing interventions targeting hypothalamic dysfunction after estrogen withdrawal.

## Results

### Progressive endocrine and thermoregulatory changes after ovariectomy in mice

To model the time-dependent effects of estrogen withdrawal on the hypothalamus, we assessed a series of timepoints post-OVX in wild-type female mice (**Fig. 1a**). Serum luteinizing hormone (LH) concentrations increased progressively, with levels peaking at 2 months post-OVX before declining at 4 months post-OVX (**Fig. 1b**). This biphasic LH pattern, initial rise followed by decline, parallels the endocrine trajectory observed during human reproductive aging, in which gonadotropin levels increase during perimenopause and subsequently decline in late postmenopause^19^. To characterize changes in thermoregulation, core temperature was measured across timepoints, revealing a peak at 14 days post-OVX that progressively normalized by 4 months (**Fig. 1c**). Together, these physiological data indicate that the 14-day post-OVX model captures the acute neuroendocrine response to estrogen withdrawal, characterized by elevated gonadotropins and disrupted thermoregulation, while the 4-month post-OVX model reflects a more chronic, adapted state with declining gonadotropins and normalized temperature.

**Figure 1.**
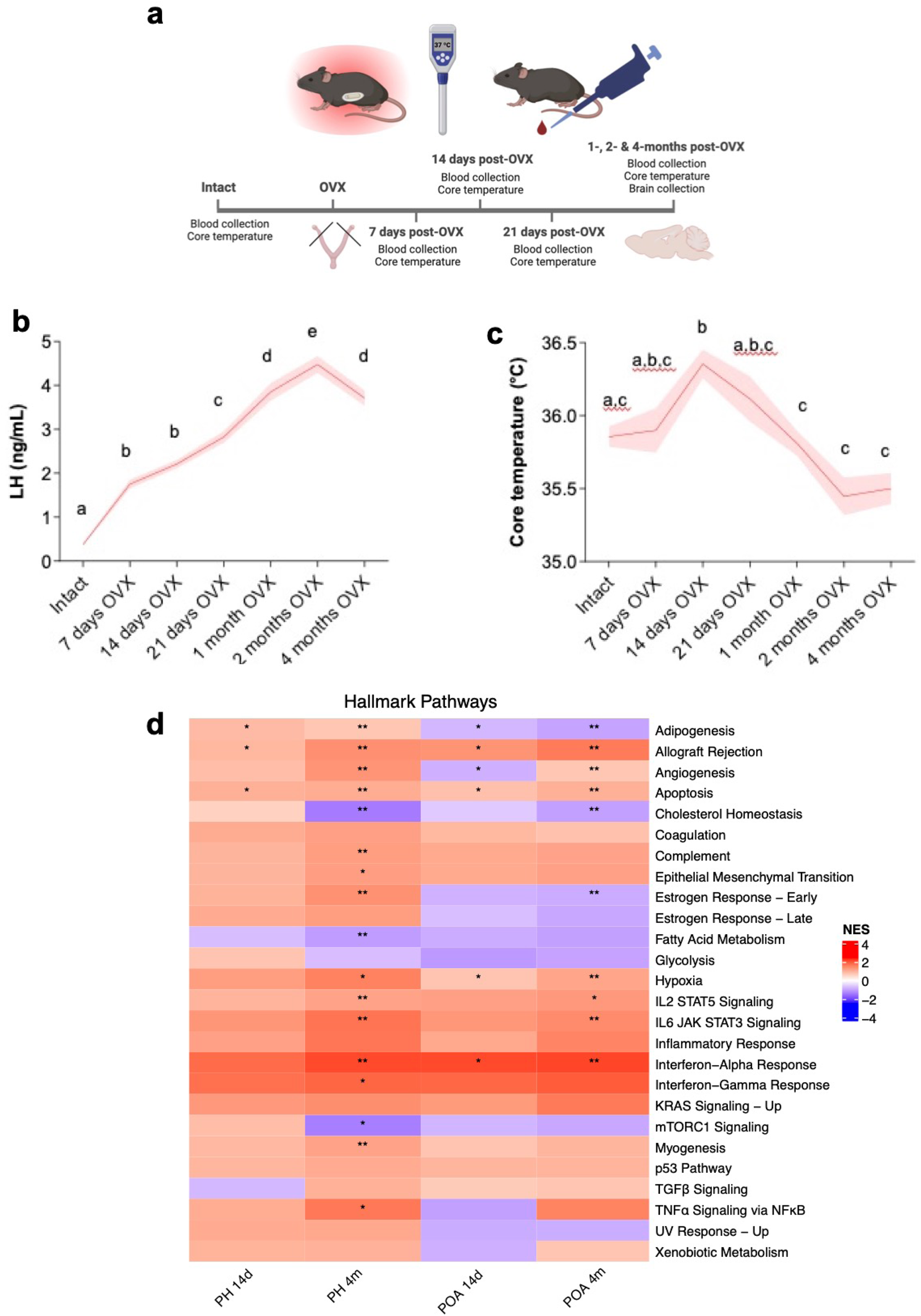
Progressive physiological and transcriptomic changes in the mouse hypothalamus after ovariectomy. a) Schematic representation of the experimental design. b) Luteinizing Hormone (LH) concentration in serum from whole blood at 7-, 14-, 21-days and 1-, 2- and 4-months post-OVX in wild-type female mice (n=20/group). c) Core temperature (°C) measurements at 7-, 14-, 21-days and 1-, 2- and 4-months post-OVX in wild-type female mice (n=20/group). b–c) Data shown as mean ± SEM. Groups with different letters are significantly different, P < 0.05 by One-way ANOVA/Tukey. d) Heatmap of normalized enrichment scores (NES) for Hallmark gene sets in the Posterior Hypothalamus (PH) and the Preoptic Area (POA); significant enrichments indicated by asterisks (* p-adj <0.05; ** p-adj < 0.01; *** p-adj < 0.001).

### Time-dependent hypothalamic transcriptomic changes after OVX

To determine whether these physiological changes are accompanied by corresponding molecular alterations in the hypothalamus, we performed bulk RNA-sequencing of intact, 14 days (short-term) post-OVX, and 4 months (long-term) post-OVX models. We selected two regions of the hypothalamus based on the localization of neurons controlling temperature regulation (preoptic area, POA) and the localization of neurons controlling metabolic and reproductive centers (posterior hypothalamus, PH). Cell type deconvolution analyses of the bulk RNA-sequencing data revealed the presence of previously reported hypothalamic cell types^20^ across all samples (**Supplementary Fig. 1**).

Global gene set enrichment analysis (GSEA)^21^ revealed marked enrichment in inflammatory pathways, with the greatest changes observed at 4 months post-OVX in the PH and POA (**Fig. 1d**, **Supplementary Table 1**). TNF-alpha signaling via NF-_κ_B, interferon responses, IL6-JAK-STAT3 signaling, and complement pathways were among the most significantly enriched. STRING analysis^22^ of TNF-alpha hallmark pathway leading edge genes highlighted genes related to thermoregulation^23^, including positive regulation of fever generation and prostaglandin biosynthesis^24^ (**Supplementary Fig. 2**). Notably, estrogen response pathways showed progressive depletion with time after OVX, particularly in the POA, consistent with the loss of ovarian estrogen signaling. These data demonstrate that the hypothalamic inflammatory response to estrogen withdrawal is not immediate but rather develops progressively, with the most robust transcriptomic changes emerging months after OVX.

### Differential gene expression reveals distinct short-term and long-term OVX signatures

We next conducted differential gene expression^25^ analyses between intact, short-term and long-term post-OVX models in both the PH and POA. Volcano plots highlight significantly differentially expressed genes (FDR < 0.05) between conditions and regions (**Fig. 2a–d**, **Supplementary Fig. 3a,b**, **Supplementary Table 2**). Notably, the number of DEGs increased substantially at 4 months relative to 14 days post-OVX. Heatmaps depicting how the identified genes are expressed across conditions in the PH (**Fig. 2e**) and POA (**Fig. 2f**) (**Supplementary Table 3**) revealed distinct temporal profiles: a subset of genes was already altered at 14 days post-OVX, while a larger set showed changes only at 4 months, and some showed progressive changes across both timepoints.

**Figure 2.**
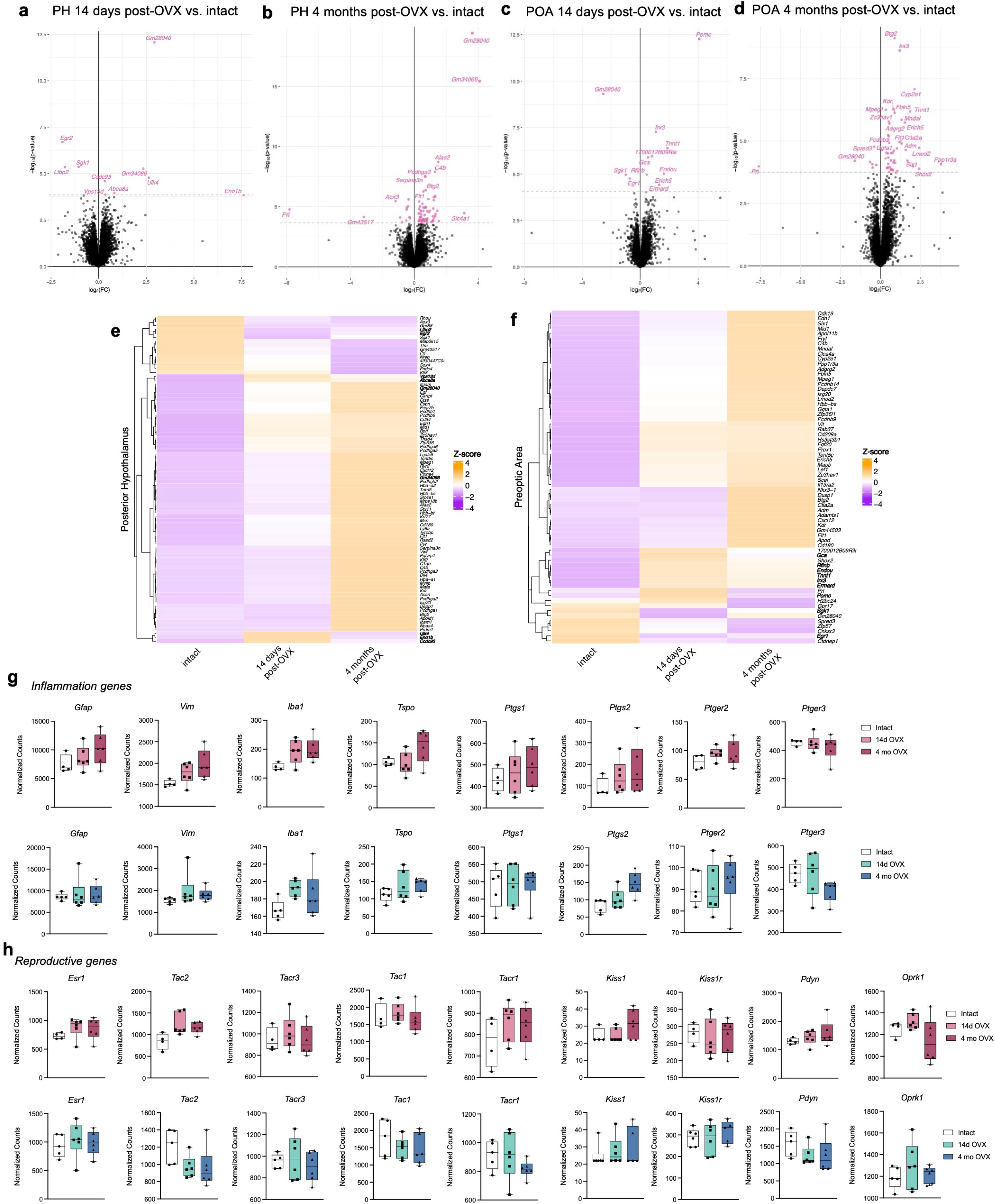
Differential gene expression and candidate gene analysis in the post-OVX mouse hypothalamus. a–d) Volcano plots highlighting significantly differentially expressed genes (FDR < 0.05, labeled in pink) between: a) PH 14 days post-OVX vs. intact (13 protein-coding DEGs), b) PH 4 months post-OVX vs. intact (82 protein-coding DEGs), c) POA 14 days post-OVX vs. intact (14 protein-coding DEGs), d) POA 4 months post-OVX vs. intact (70 protein-coding DEGs). e) Expression of PH DE genes across groups. f) Expression of POA DE genes across groups. g) Expression of inflammation-related, astrocyte and microglia marker genes in the PH (top row) and POA (bottom row). h) Expression of KNDy genes and their receptors in the PH (top row) and POA (bottom row).

Among the differentially expressed genes, *Gm28040*, which shares overlapping CDS with *Kiss1*, showed the largest expression change, increased at 14 days and 4 months OVX in the PH and decreased in both groups in the POA, resembling the known region-specific regulation of *Kiss1* in both hypothalamic areas^26^. This suggests that in the mouse, *Gm28040* encodes a kisspeptin isoform (**Fig. 2a–d**, **Supplementary Table 2**).

We then examined the expression of previously identified marker genes for inflammation (**Fig. 2g**) and reproduction (**Fig. 2h**) in both the PH and the POA. While individual KNDy and inflammation-related genes did not reach statistical significance after adjustment for multiple hypothesis testing, the expression of inflammation and astrocytic markers showed increasing trends from intact to 4 months post-OVX, with notable increases in *Vim*, *Iba1*, and *Tspo* in the PH, and in the prostaglandin pathway (*Ptgs2*) in the POA. The KNDy genes, involved in reproductive and thermoregulatory (VMS) control, showed the expected increase in expression in the PH (where the arcuate nucleus is located) in proportion to time after OVX, consistent with progressive loss of estrogen negative feedback on these sex-steroid-regulated genes^27^.

### Protein-level validation of KNDy neuron changes and neuroinflammation

To validate key transcriptomic findings at the protein level, we performed immunofluorescence staining for neurokinin B (NKB), the early activation marker c-Fos, and the astrocyte marker GFAP in the arcuate nucleus (**Fig. 3**). NKB immunoreactivity was significantly reduced in both 14-day and 4-month post-OVX mice compared to intact controls (**Fig. 3a,d,g, j**). This decrease in NKB peptide stores, in the context of increased *Tac2* mRNA expression (**Fig. 2h**), is consistent with enhanced neuropeptide release and depletion rather than decreased synthesis, a hallmark of hyperactivated neuropeptide systems.

**Figure 3.**
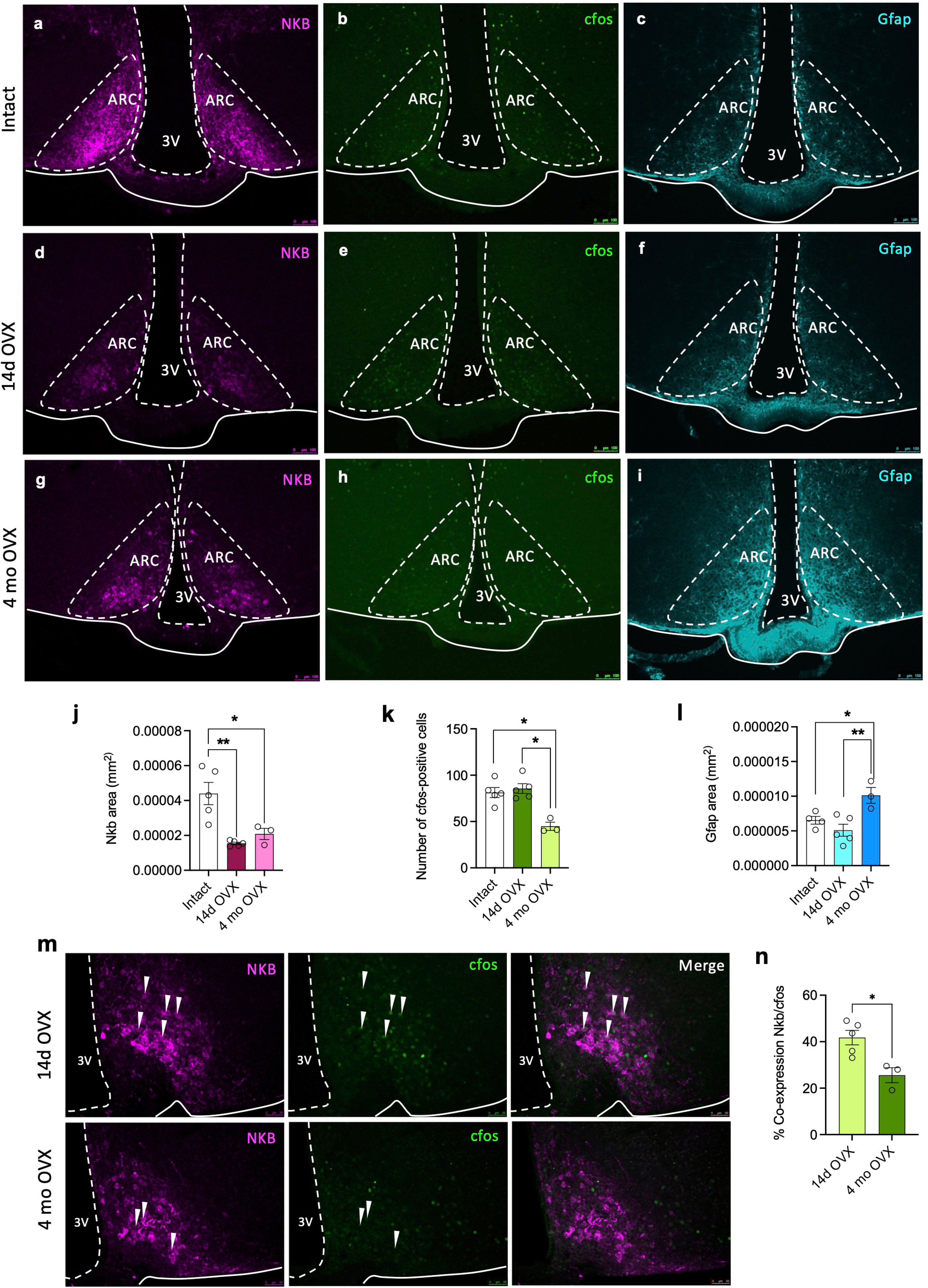
Differential neurokinin B distribution, neuronal activity and astrocyte reactivity in the arcuate nucleus after ovariectomy. Representative immunofluorescence staining of NKB (a, d, g), cFos (b, e, h) and GFAP (c, f, i) in the arcuate nucleus (ARC) from intact (a–c), 14 days post-OVX (d–f), and 4 months post-OVX female mice (g–i). j) Quantification of NKB area (mm²) in the ARC from intact, 14 days post- and 4 months post-OVX female mice. k) Quantification of cFos-positive cells in the ARC from intact, 14 days post- and 4 months post-OVX female mice. l) Quantification of GFAP area (mm²) in the ARC from intact, 14 days post-and 4 months post-OVX female mice. m) Representative images of NKB and cFos co-localization in the ARC from 14 days and 4 months post-OVX female mice. n) Percentage of NKB/cFos co-expression. Data from j–l and n shown as mean ± SEM. Statistical significance determined by Mann-Whitney test (intact vs. OVX). *P < 0.05, **P < 0.01.

c-Fos expression in the arcuate nucleus remained elevated at 14 days post-OVX at a similar level to control gonad intact mice and declined significantly by 4 months (**Fig. 3b,e,h, k**). To determine whether this decline in activity occurred specifically in KNDy neurons, we quantified NKB/c-Fos co-localization, which revealed a ∼50% decrease in the proportion of activated KNDy neurons at 4 months compared to 14 days post-OVX (**Fig. 3m, n**). This reduction in KNDy neuron activity at long-term post-OVX aligns with the normalization of core body temperature observed at this timepoint (**Fig. 1c**) and provides a cellular basis for the temporal dynamics of vasomotor-like responses.

GFAP immunoreactivity increased progressively with time after OVX, with significant elevations at both 14 days and 4 months compared to intact animals (**Fig. 3c,f,i, l**). This increase in astrocytic reactivity corroborates the transcriptomic evidence for progressive neuroinflammation and glial activation in the hypothalamus following estrogen withdrawal.

### Age-related transcriptomic changes in the human female hypothalamus

Having established the temporal dynamics of hypothalamic changes after experimental estrogen withdrawal in mice, we next sought to determine whether analogous molecular signatures could be identified in the human hypothalamus. We leveraged publicly available bulk RNA-sequencing data from the Genotype-Tissue Expression (GTEx) project^28^ to examine hypothalamic gene expression changes across the adult female lifespan. It is important to note that this human dataset, while the best currently available resource, has an inherent limitation since samples derive from postmortem tissue of 31 female donors (ages 28–65) whose menopausal status and hormone therapy history are unavailable. We therefore present these findings as age-related transcriptomic patterns in the human hypothalamus, with the caveat that associations with menopausal status, while biologically plausible given the known timing of menopause, cannot be definitively established from this dataset alone.

Female donors with hypothalamic tissue were included if they met GTEx donor eligibility criteria and their sequenced tissue had an RNA integrity number of 6 or greater (**Supplementary Table 4**). The 31 samples span the adult female age range (**Fig. 4a,b**). Surrogate variable analysis (SVA)^29^ was performed on the RNA expression matrix prior to gene expression analyses to control for latent, confounding sources of variation in the data, such as postmortem interval, donor cause of death, and hormone therapy status. Cell type deconvolution of the bulk RNA-sequencing data revealed the presence of previously reported hypothalamic cell types^20^ (**Supplementary Fig. 4a**, **Supplementary Table 5**) and the presence of all eight major transcription factors (*FEZF1, FOXB1, LHX6, MEIS2, OTP, SIM1, SIX3, TBX3*) identified in the human hypothalamus from the HYPOMAP single-nucleus RNA-seq reference atlas^30^ (**Supplementary Fig. 4b**), indicating that there are no major differences in cell type composition between samples in the cohort.

**Figure 4.**
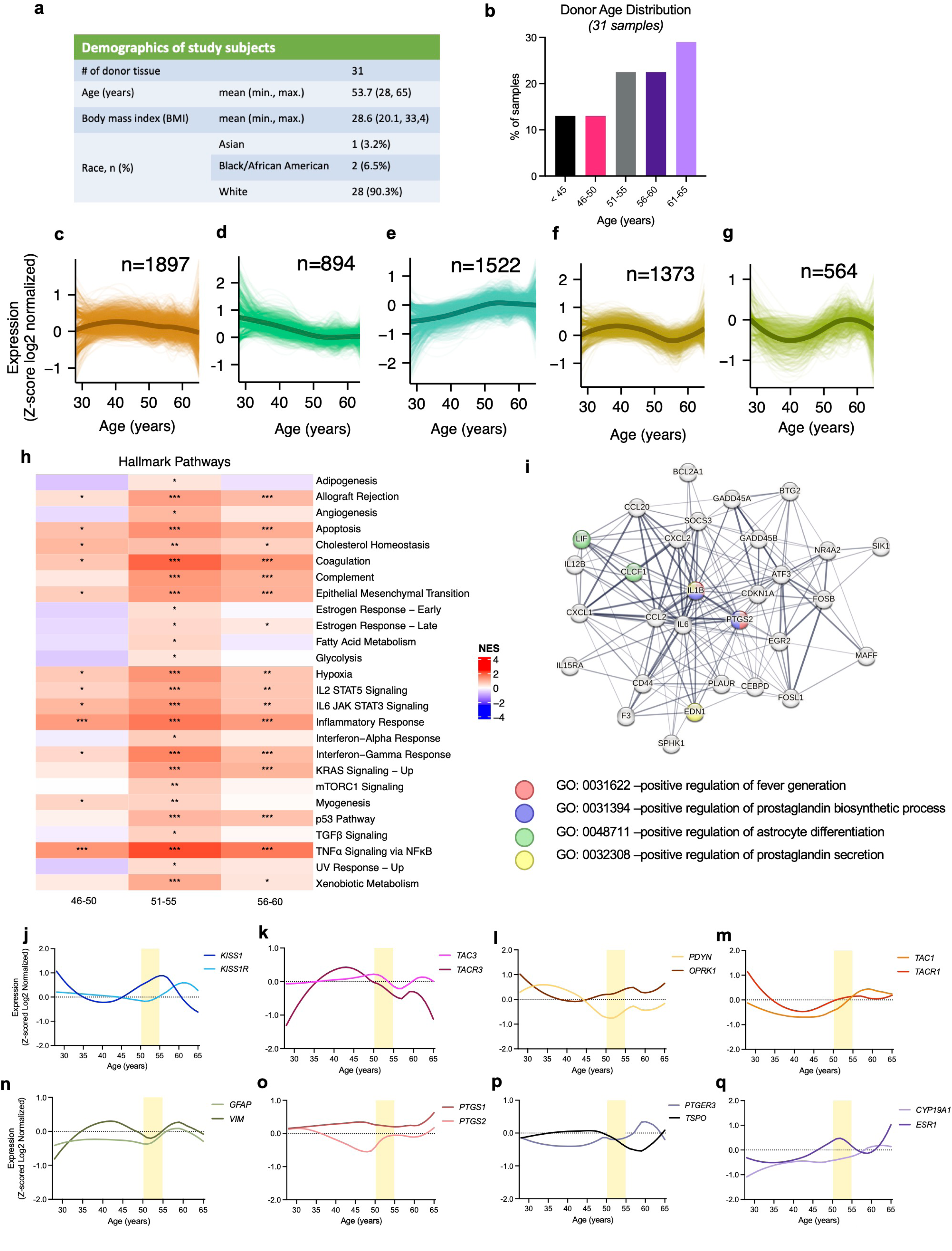
Gene expression trajectories and pathway analysis in the human female hypothalamus across chronological age. a) Demographic overview of the GTEx donor subset used in this study. b) Distribution of ages within the donor cohort and the defined age groupings for subsequent analyses. c–g) Sets of genes grouped by hierarchical clustering and displayed using LOESS trajectories exhibit distinct patterns: c) constant expression across age, d) decreased expression across age, e) increased expression across age, f) decreased expression during ages 51–60, g) increased expression during ages 51–60. h) Heatmap of normalized enrichment scores (NES) for Hallmark gene sets in which there was a statistically significant pathway enrichment in at least one of the three age-group comparisons (46–50, 51–55, and 56–60 vs. <45); significant enrichments indicated by asterisks (* p-adj <0.05; ** p-adj < 0.01; *** p-adj < 0.001). i) STRING analysis of TNF-alpha hallmark pathway leading edge genes between the 51–55 and <45 age groups. j–q) LOESS trajectories of KNDy genes and their receptors, thermoregulatory genes, astrocyte- and microglia-marker genes across age in human hypothalamus tissue: j) *KISS1* and *KISS1R*, k) *TAC3* and *TACR3*, l) *PDYN* and *OPRK1*, m) *TAC1* and *TACR1*, n) *GFAP* and *VIM*, o) *PTGS1* and *PTGS2*, p) *PTGER3* and *TSPO*, q) *CYP19A1* and *ESR1*.

We assessed patterns of gene expression changes across age using locally estimated scatterplot smoothing (LOESS). LOESS expression profiles were grouped by hierarchical clustering, revealing distinct patterns, including genes with nearly constant (**Fig. 4c**), decreased (**Fig. 4d**), and increased (**Fig. 4e**) expression across age. Of particular interest was the identification of gene clusters with specific decreases (**Fig. 4f**) or increases (**Fig. 4g**) coinciding with the age range during which menopause typically occurs (∼45–55 years) (**Supplementary Table 6**), though we emphasize that without direct menopausal status data, we cannot attribute these changes solely to menopause rather than to other age-related processes.

Next, we divided the cohort into age bins informed by epidemiological data from the Study of Women’s Health Across the Nation (SWAN), where the median age at final menstrual period is 51–52 years^31^ (**Fig. 4b**): <45 years (n=4), 46–50 years (n=4), 51–55 years (n=7), 56–60 years (n=7), and 61–65 years (n=9). We acknowledge that individual variation in menopausal timing means these bins represent chronological age ranges that statistically correspond to typical reproductive stages in population studies, such as SWAN, but we cannot assign clinical menopausal staging to individual donors. Gene set enrichment analysis (GSEA)^21^ between each age group and the <45 group revealed the highest and most significant enrichments in inflammatory pathway signaling in the 51–55 age group (**Fig. 4h**, **Supplementary Table 7**). These include TNF-alpha signaling via NF-_κ_B, interferon responses, IL2-STAT5 and IL6-JAK-STAT3 signaling, and complement pathways, strikingly similar to the pathways enriched in the long-term OVX mouse model (**Fig. 1d**). STRING analysis^22^ of TNF-alpha hallmark pathway leading edge genes between the 51–55 and <45 age groups highlighted genes involved in thermoregulation^23,24^, including fever responses (**Fig. 4i**).

We examined the LOESS trajectories of key ligand-receptor systems related to KNDy signaling, thermoregulation, and glial activation across age in the human hypothalamus (**Fig. 4j–q**). *KISS1* and *ESR1* showed marked increases during the typical menopausal years, consistent with loss of estrogen negative feedback^32,33^, followed by a decline ∼10 years after their peak. *TACR3* expression increased in the years leading up to age 50 and then progressively declined. The astrocyte activation markers *GFAP* and *VIM* increased during the 50–55 age range, while the microglial marker *TSPO* increased in later years, as previously reported^34^. Changes in prostaglandin synthases (*PTGS1/2*) and receptors (*PTGER3*) were also observed, with *PTGER3* showing a notable increase followed by a sharp decrease around age 60. We additionally assessed the expression of *PGR*, *GPER1*, and *ESR2* (**Supplementary Fig. 5**) based on a recent publication examining these genes as potential postmortem biomarkers in the hypothalamus^35^.

### Identification of age-associated differentially expressed genes in the human hypothalamus

We performed differential gene expression analysis between the 51–55 and <45 age groups (**Fig. 5a**, **Supplementary Table 8**). Twenty-eight protein-coding genes reached significance at FDR < 0.05, with the majority showing increased expression in the older age group. We then examined the expression of these differentially expressed genes across all defined age groups (**Fig. 5b**, **Supplementary Table 9**), revealing that many of the identified genes show progressive changes with age rather than abrupt shifts at a single transition point. The two genes with the most significant adjusted p-values were *AKAP5* (A-kinase anchoring protein 5), which decreased with age, and *CDKN1A* (p21), which increased (**Fig. 5c, d**). We note that these genes, while identified as DEGs in the human dataset, did not reach significance in the mouse OVX model, which may reflect species-specific regulation, differences in statistical power between the datasets, or the distinct nature of surgical versus natural estrogen withdrawal. Spatial transcriptomic visualization of *Akap5* and *Cdkn1a* from the Allen Brain Atlas confirmed their expression in hypothalamic subregions including the DMH, VMH, and ARH (**Supplementary Fig. 6**). Additionally, we assessed expression of canonical aging-associated genes^36^ across our groups (**Supplementary Fig. 7a**) and plotted trajectories for genes with previously reported functions in brain aging and cognitive decline^37–41^ (**Supplementary Fig. 7b**).

**Figure 5.**
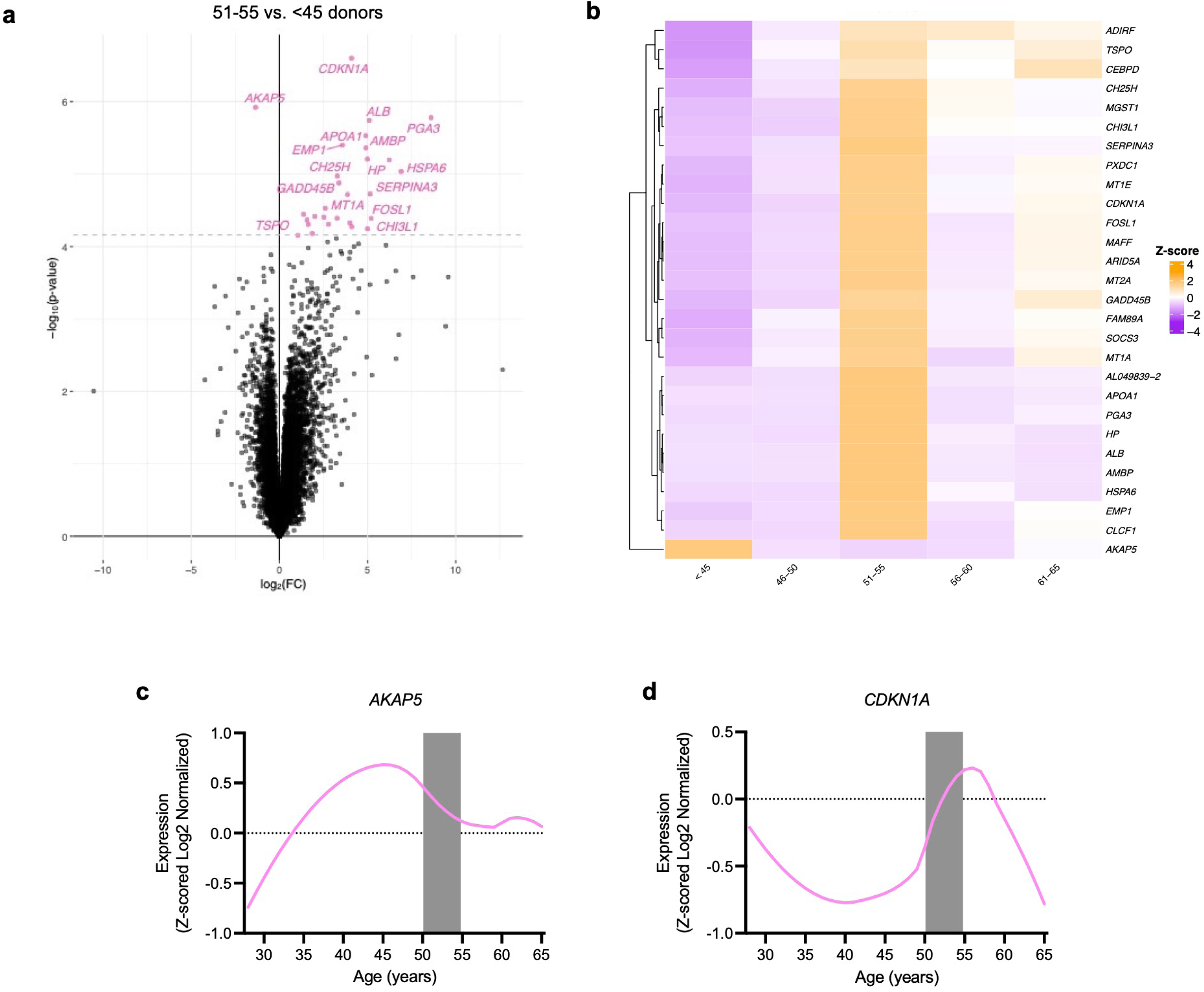
Age-associated differentially expressed genes in the human hypothalamus. a) Volcano plot highlighting significantly differentially expressed genes between the 51–55 and <45 age groups. Genes with FDR < 0.05 are highlighted in pink. b) Expression of genes with FDR < 0.05 across age groups. c) LOESS trajectory of *AKAP5* across age. d) LOESS trajectory of *CDKN1A* across age.

### Cross-species comparison reveals conserved hypothalamic inflammatory signatures

A substantial proportion of expressed genes are shared between the human and mouse hypothalamus (∼77%; **Supplementary Fig. 8**). To assess whether the transcriptomic changes observed in the mouse OVX model capture similar changes to those identified in the human hypothalamus across age, we compared GSEA results between the two datasets. This analysis revealed statistically significant pathway enrichment correlations (**Fig. 6a**), with the strongest correlations observed between the 4-month post-OVX PH and the 51–55 age group in humans. This supports the proposition that the long-term OVX model captures core transcriptomic features that parallel age-related changes in the human female hypothalamus, though we recognize that the human changes may reflect both estrogen loss and other age-related processes.

**Figure 6.**
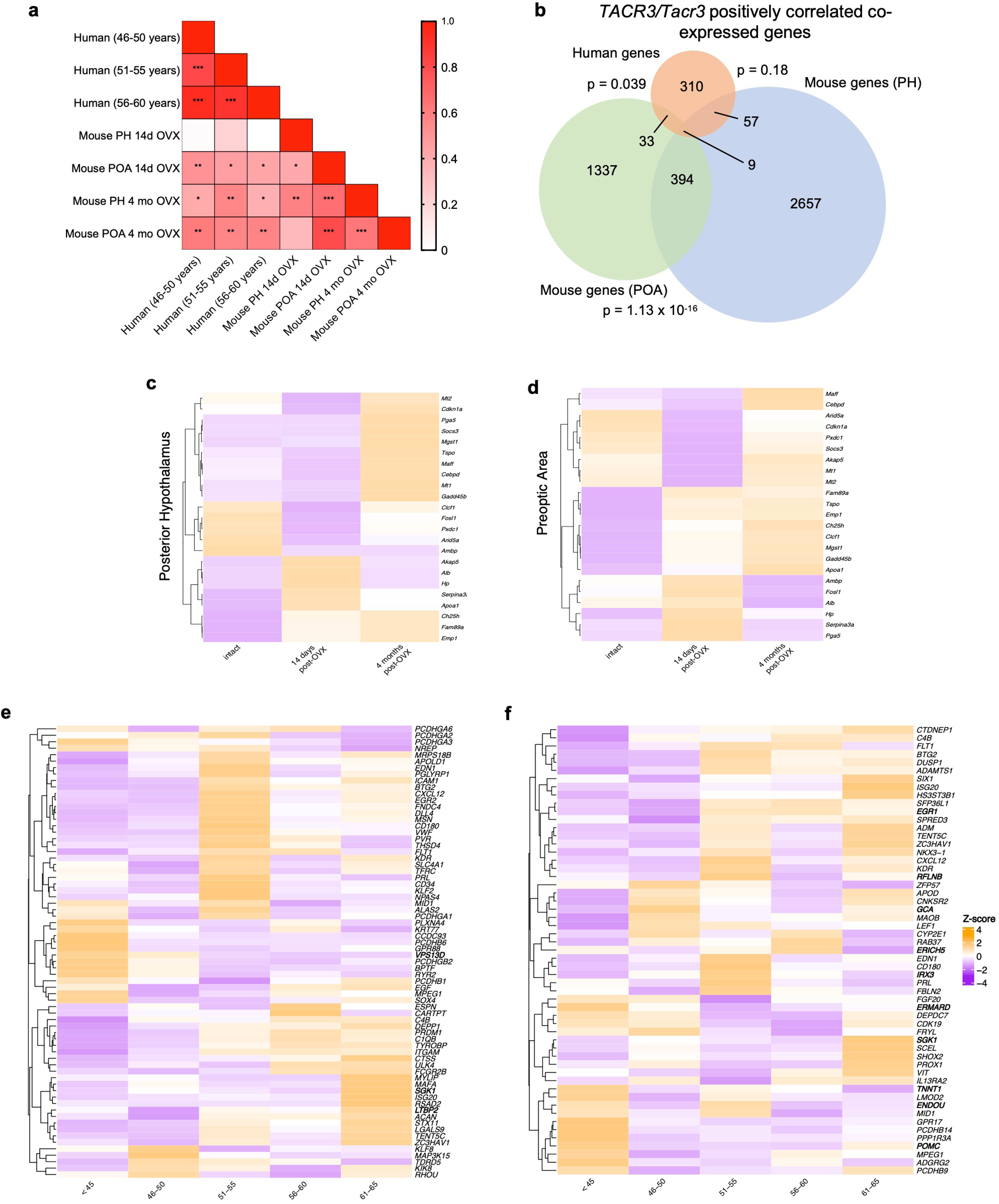
Cross-species comparison of hypothalamic transcriptomic changes. a) Heatmap of GSEA pathway correlations between human age groups and mouse OVX conditions; p-value from Pearson’s correlation. b) Overlap of *TACR3/Tacr3* positively correlated co-expressed genes between human and mouse; p-value from hypergeometric test. c–d) Expression of identified DE genes from the human dataset across conditions in the mouse c) PH and d) POA. e) Expression of identified DE genes from the mouse PH across age groups in the human dataset. f) Expression of identified DE genes from the mouse POA across age groups in the human dataset.

We also compared GSEA results between our OVX mouse model and a published single-cell RNA-seq dataset of the aging female mouse hypothalamus^20^ to assess whether the OVX model captures similar hypothalamic changes to those associated with natural aging in mice. Of the 26 Hallmark pathways significantly impacted in the human dataset (**Supplementary Table 7**), only eight were similarly altered in the aged mouse hypothalamus^20^, compared with 14 in the OVX model. Notably, the aged mouse study^20^ did not report any changes in the estrogen response or TNF_α_ signaling pathways, which are consistently affected in both the human dataset and the OVX mouse model and are directly relevant to reproductive aging^42^. At the cell-type level, the observed pathway changes in macrophages and microglia from the aged mouse dataset exhibited the greatest similarity to those changes observed in the human and OVX mouse model (**Supplementary Table 10**). Despite this partial overlap, the two mouse models share a single significantly differentially expressed gene (*Thsd4*), suggesting that they capture largely distinct hypothalamic transcriptional programs, the OVX model preferentially modeling estrogen-dependent changes and the aging model capturing broader senescence-related alterations.

To determine whether cross-species similarities extend beyond pathway-level analyses, we selected *TACR3*, given its documented role in VMS onset and severity^43^, and performed co-expression analyses. *TACR3/Tacr3* co-expression analysis identified genes whose expression positively correlates with *TACR3* in human and *Tacr3* in the OVX mouse POA revealing statistically significant overlap in the positively co-regulated gene sets (hypergeometric test, p = 0.039; **Fig. 6b**, **Supplementary Table 11**). We note that *TACR3/Tacr3* itself was not identified as a DEG in either species in the pairwise comparisons, but that its co-expression network is conserved, suggesting that the broader transcriptional program in which it participates is similarly affected by estrogen loss across species.

Finally, we examined whether DE genes identified in one species showed concordant changes in the other. When human DEGs were examined across mouse conditions, two-thirds of the genes differentially expressed in the 51–55 age group showed a clear increase at 4 months post-OVX, while the remaining third were captured at 14 days post-OVX, in both PH and POA (**Fig. 6c, d**). Conversely, the majority of mouse PH and POA DEGs that showed the highest changes at 4 months post-OVX were upregulated in the 51–55 age group (**Fig. 6e, f**), though individual genes such as *AKAP5* and *CDKN1A* (identified as human DEGs) were not differentially expressed in the mouse, underscoring the expected species-specific differences alongside the broader pathway-level conservation.

## Discussion

This study provides the first systematic temporal characterization of hypothalamic transcriptomic changes following estrogen withdrawal, revealing progressive neuroinflammation that develops over months in the mouse and whose molecular characteristics parallel age-related changes in the female human hypothalamus.

A central finding is that the hypothalamic response to estrogen loss is markedly time-dependent. While physiological changes (i.e. elevated LH and core temperature) manifest rapidly after OVX, the full transcriptomic inflammatory signature does not emerge until 4 months post-OVX. This dissociation between acute neuroendocrine responses and delayed transcriptomic remodeling has important implications since short-term OVX studies, which constitute the majority of the literature, may capture the initial endocrine disruption but miss the progressive neuroinflammatory processes that could underlie the chronic sequelae of estrogen withdrawal. The delayed onset of robust inflammatory pathway activation, particularly TNF-alpha signaling via NF-_κ_B and interferon responses, suggests that neuroinflammation is not a direct consequence of estrogen loss but rather develops as a secondary process, possibly driven by cumulative glial activation and loss of estrogen’s anti-inflammatory protective effects.

Our protein-level validation in the arcuate nucleus provides cellular resolution to complement the transcriptomic data. The finding that NKB peptide stores are depleted while *Tac2* mRNA is increased is consistent with a state of hyperactivated neuropeptide release in KNDy neurons, particularly at 14 days post-OVX. The subsequent ∼50% decline in KNDy neuron activation (c-Fos/NKB co-localization) at 4 months aligns with the normalization of core temperature at this timepoint and offers a potential cellular mechanism for the resolution of VMS-like thermoregulatory disruption. This temporal pattern is consistent with clinical observations that hot flashes, while beginning during perimenopause, typically resolve within 5–10 years after menopause^18^, and may reflect adaptive processes within KNDy neurons to prolonged estrogen deficiency. In parallel, the progressive increase in GFAP immunoreactivity demonstrates that astrocytic reactivity continues to build even as acute neuronal hyperactivation resolves, pointing to distinct temporal dynamics for neuronal versus glial responses to estrogen withdrawal.

The concurrent increase in inflammatory signaling and astrocyte/microglia markers has functional implications beyond the hypothalamus. Activated astrocytes and microglia in thermoregulatory and metabolic hypothalamic nuclei can modulate local neuronal excitability through altered neurotransmitter reuptake, gap junction coupling, or cytokine release^44^. The prostaglandin pathway changes we observed directly link to autonomic output controlling cutaneous vasodilation, the effector mechanism of VMS^45,46^. Furthermore, hypothalamic inflammation can influence systemic metabolic regulation through altered brown adipose tissue thermogenesis, potentially contributing to weight gain commonly observed after menopause^47^. The upregulation of NF-_κ_B-mediated inflammatory signaling may also propagate to peripheral tissues through neuroendocrine and autonomic pathways, contributing to the systemic low-grade inflammation (“inflammaging”) associated with cardiometabolic risk in postmenopausal women^48^.

To place our experimental findings in a translational context, we analyzed human hypothalamic transcriptomic data from the GTEx consortium. While this observational dataset has important limitations based on the modest cohort size (n=31), reliance on chronological age as a proxy for menopausal status, use of bulk RNA-seq in a heterogeneous tissue, and inability to control for hormone therapy or menopausal symptoms, it represents the most comprehensive publicly available resource for studying human hypothalamic gene expression across the female lifespan. Thus, we explicitly framed the human data as age-related transcriptomic changes rather than menopausal changes.

Despite these caveats, the convergence between mouse and human datasets at the pathway level is notable. The inflammatory pathway enrichments observed in the 51–55 age group closely mirror those seen at 4 months post-OVX, and GSEA correlation analyses confirmed significant pathway-level similarity between the long-term OVX mouse PH and the human hypothalamus during the age range typically corresponding to late perimenopause and early post-menopause. That two-thirds of differentially expressed genes in the human dataset showed concordant changes at 4 months post-OVX, rather than at 14 days, further supports the idea that the long-term OVX model preferentially captures the transcriptomic state associated with the later stages of reproductive aging, while the short-term model may better represent the acute peri-menopausal phase.

The human data also revealed genes of interest, including *AKAP5* and *CDKN1A* (p21), as the most significantly differentially expressed between the 51–55 and <45 age groups. *AKAP5* encodes a scaffolding protein involved in synaptic plasticity^49–52^ and showed decreased expression with age in the hypothalamus. *CDKN1A*, encoding the cell cycle regulator p21^53^, was upregulated and may promote senescence-associated secretory phenotype (SASP)^53–56^, potentially contributing to the inflammatory phenotype observed. Notably, these genes were not differentially expressed in the mouse OVX model, which could reflect species-specific regulation, differences in the nature of surgical versus natural estrogen withdrawal, or the contributions of aging-related processes beyond estrogen loss in the human dataset. These observations highlight the complexity of disentangling estrogen-dependent from age-dependent changes and underscore the value of using multiple complementary approaches.

Our analysis of KNDy gene expression in the human hypothalamus, while interpreted with caution given the constraints of the dataset, revealed patterns consistent with known biology. *KISS1* expression appeared to increase with age during the typical menopausal years, consistent with reports of kisspeptin neuron hypertrophy in postmenopausal women^32,33^, followed by a decline. *TACR3* expression showed a distinct trajectory, increasing before the typical age of menopause and declining thereafter, which aligns with the known time course of VMS^18^. While these KNDy genes did not reach significance as DEGs after correction for multiple testing, likely reflecting the limited sample size, their expression trajectories are biologically consistent with the established literature and with our experimental findings in the mouse model. Thus, our data offer novel insights into the mechanisms that could mediate the decrease in LH levels and termination of hot flashes years after menopause, likely mediated by the significant drop in hypothalamic *KISS1* and *TACR3* expression^57,58^.

Regarding the choice of animal model, while multiple rodent models exist to study the consequences of estrogen loss, we selected long-term OVX because it provides precise control over the timing of estrogen withdrawal, enabling direct temporal alignment of endocrine, thermoregulatory, and transcriptomic changes. The VCD (4-vinylcyclohexene diepoxide) model offers an alternative approach that more closely mimics gradual follicular depletion^59^, but introduces confounders including variable timing of ovarian failure (15–160 days), residual androgen production^59–61^, and potential off-target effects^62,63^. Our OVX model, while representing abrupt surgical menopause, provides a robust framework for studying chronic hypoestrogenism and enables the kind of precise temporal analyses we describe here.

In conclusion, this study provides a temporal framework for understanding how estrogen withdrawal progressively reshapes the hypothalamic transcriptome, moving from acute neuroendocrine disruption to chronic neuroinflammation. The convergent inflammatory signatures observed across species suggest that this neuroinflammatory program is a conserved consequence of estrogen loss, offering both mechanistic insight and a preclinical platform for testing interventions. Future studies using single-cell approaches in both species will be essential to resolve cell-type-specific contributions and establish causal relationships between these transcriptomic changes and menopausal symptoms.

## Materials and Methods

### Mouse studies

#### Study approval and animals

The animal studies were approved by the Brigham and Women’s Hospital Institutional Animal Care and Use Committee (IACUC) in the Center for Comparative Medicine. Adult wild-type (WT) C57/BL6 female mice were group housed under constant conditions of temperature (22–24°C) and light (12:12-hour light:dark cycle), fed with standard mouse chow and ad libitum access to tap water.

#### Experimental design

To investigate the effects of sex steroid withdrawal on LH levels, core temperature (Tc), and hypothalamic transcriptome profile, intact WT female mice were bilaterally ovariectomized (OVX) to generate models at progressive timepoints after estrogen withdrawal. Blood LH levels and Tc were measured at baseline (intact), at 7-, 14- and 21-days post-OVX and at 1-, 2- and 4-months post-OVX. Additional females were euthanized before OVX in diestrus (intact group), after 14 days of OVX (short-term post-OVX group) or after 4 months of OVX (long-term post-OVX group) and their posterior (PH) and preoptic area (POA) of the hypothalamus were collected for transcriptome profile analysis by bulk RNA-seq.

#### Ovariectomy (OVX)

Adult (3–4 months-old) female mice were bilaterally OVX under isoflurane anesthesia. Mice were treated with Buprenex SR (0.1 mg/kg, SC) and meloxicam (5 mg/kg, SC) on the day of surgery. Mice were allowed to recover for at least 1 week before the start of experiments.

#### Whole blood samples and LH ELISA

For LH measurements, a 4µl blood sample was collected with a pipette. Whole blood was immediately diluted in 116µl of 0.05% PBST [phosphate buffer saline (Boston Bio Products, catalog #BM220) containing Tween-20 (Sigma, catalog #P2287)], vortexed, and frozen on dry ice. Samples were stored at −80°C until analyzed using an in-house sandwich enzyme-linked immunosorbent assay as previously described^64^.

#### Core temperature (Tc)

For Tc measurements, intact female mice were implanted (intraperitoneally) with an IPTT-300 temperature transponder (BioMedic Data Systems, Inc.). Tc from the transponder were collected with a DAS-8007 Reader and DASHost 8000 software (BioMedic Data Systems, Inc.).

#### Immunohistochemistry

Animals were anesthetized with a ketamine/xylazine/saline cocktail and transcardially perfused with saline (0.9% NaCl) followed by 4% paraformaldehyde (PFA) diluted in phosphate buffer (PB) (Boston BioProducts, Inc). Brains were removed, stored in 4% PFA overnight and then transferred into sucrose solution (30% sucrose in 0.1 M PB, Thermo Fisher Scientific) at 4°C. After sucrose infiltration, the tissue was frozen in Tissue-Tek O.C.T. Compound (Sakura Finetek-4583) and cut on a freezing stage microtome (Microm HM 450, Thermo Fisher Scientific) into 30 µm coronal sections. Free-floating sections were washed in tris buffered saline (TBS, Boston BioProducts, Inc) and blocked for 1 hour at room temperature in incubation solution [TBS + 0.3% Triton X-100 (Millipore Sigma) + 2% normal goat serum (NGS)]. Sections were subsequently incubated for 48 hours at 4°C with primary antibodies: rabbit-anti neurokinin B (1:1000, Novus Biologicals, catalog #NB300-201), guinea pig anti-cFos (1:1000, Synaptic System, catalog #226308) and mouse anti-GFAP (1:300, Cell Signaling, catalog #3670) diluted in incubation solution. Next, sections were incubated for 1 hour at room temperature with secondary antibodies: anti-rabbit Alexa Fluor 594 (1:500, Thermo Fisher Scientific, catalog #A11012), anti-guinea pig Alexa Fluor 488 (1:500, Jackson ImmunoResearch, catalog #706545148) and anti-mouse donkey Alexa 594 (1:500, Invitrogen, catalog #A32744). Sections were mounted onto Super-frost plus glass slides (Fisher Laboratories), air dried and cover slipped with Vectashield mounting medium with DAPI (Vector laboratories).

#### Microscopy and image analysis

The images were captured using a fluorescent microscope (Leica DM 2500) in JPG format and processed using the open-source software ImageJ, Fiji (National Institutes of Health, Bethesda, Maryland). For image quantification, representative images of the areas of interest (i.e., ARC) were delimited using as reference the mouse section from Allen Brain Atlas^65^. Both sides of bilateral ARC were counted on a similar number of sections and animal, and replicate values were averaged.

#### RNA extraction, library preparation, and sequencing

Mouse posterior hypothalamus and preoptic area regions were dissected and flash frozen prior to extracting RNA using a Zymo Quick RNA microprep kit according to the manufacturer’s instructions. RNA concentration was measured using a Qubit 4.0 fluorometer and the Qubit RNA HS Assay Kit (ThermoFisher). All samples had RNA integrity numbers (RIN) greater than 7 as measured on a Bioanalyzer 2100 instrument (Agilent). RNA sequencing libraries were prepared using the KAPA mRNA HyperPrep–Stranded Kit (Roche) and sequenced on a NovaSeq SP (Illumina) with 50bp paired-end reads.

#### RNA-seq data processing and analysis

We performed all analyses using mouse genome build mm39. To pseudoalign reads to the transcriptome annotation and estimate expression levels, we used kallisto^66^ in the tximport package^67^ (v1.24.0). We imported the resulting count data into R for analysis in DESeq2^25^ (v1.36.0). To adjust for unknown, unmodeled or latent sources of noise, we performed surrogate variable analysis (SVA) using the smartSVA package^29^ (v0.1.3) and used SV1 as a covariate in the differential gene expression analysis. Pseudoalignment and DESeq2 were performed on the entire transcriptome annotation, with downstream analysis in some cases restricted to protein-coding genes as annotated in GENCODE version M30^68^. GSEA was performed on pre-ranked ordered gene lists using the fgsea package^21^ (v1.22.0).

#### Prediction of protein interactions

The first 30 leading edge genes from the comparison of the POA between female gonadal intact vs. 4-month post-OVX mice were selected as the target gene list for TNF-alpha signaling through NF-_κ_B. The gene list was imported into STRING^22^ for network reconstruction, visualization and enrichment analysis. Medium confidence was selected as the minimum required interaction score (0.400). Enrichment biological processes defined by gene ontology (GO) were assessed.

### Human studies

#### Human hypothalamus inclusion criteria

Bulk RNA-sequencing read count data from human hypothalamus tissue was obtained from publicly available Genotype-Tissue Expression (GTEx) Project^28^ data. Donor tissue was included if: 1) donor eligibility criteria (INCEXC) == TRUE, 2) tissue RNA quality score (RIN) >= 6, 3) donors were female (SEX == 2).

#### Sex and Gender Reporting

All human hypothalamic samples from the GTEx database were from donors recorded as female (biological sex). This study examines transcriptomic changes across the female reproductive lifespan. Sex was determined by GTEx based on donor records. We use the term “women” to reflect the clinical population most directly affected by natural menopause, while acknowledging that individuals of diverse gender identities with intact ovarian function may also experience menopausal transition.

#### Human data analysis

After sub-setting the GTEx RNA expression file for our samples, we adjusted for latent sources of noise using SVA with the smartSVA package^29^ (v0.1.3). Differential expression analyses across age bins were performed using DESeq2^25^ (v1.36.0) and GSEA was performed on pre-ranked ordered gene lists using the fgsea package^21^ (v1.22.0). Gene expression trajectories across age were performed on protein-coding genes using LOESS similarly to Piehl et al. 2022^69^. Gene level output was subjected to expression trajectory correlation analysis; genes were ranked based on their correlation with *TACR3/Tacr3* expression pattern. STRING interaction analyses were performed as described above.

Given the unavailability of hormone measurements in GTEx, we used chronological age bins as proxies, informed by epidemiological data from the Study of Women’s Health Across the Nation (SWAN) where median age at final menstrual period is 51–52 years^31^. Samples were grouped into five age ranges: <45 years, 46–50 years, 51–55 years, 56–60 years, and 61–65 years. We acknowledge that individual variation in menopausal timing means our age-based classification represents chronological categories, and we cannot definitively assign clinical menopausal staging or vasomotor symptom status to individual donors. The GTEx consortium did not collect serum samples, precluding direct hormone measurements.

#### Cell type decomposition analysis

Cell type decomposition analyses to estimate cell type composition of bulk RNA-seq datasets used the BisqueRNA package^70^ (v1.0.5). Supplementary Table 5 lists the marker genes used to annotate cell types^20^.

#### Quantification and statistical analyses

Statistical tests are indicated in methods, figure legends, or text. All statistics were calculated using R software, version 4.2.1 unless stated otherwise. Differential gene expression used DESeq2 with Benjamini-Hochberg FDR correction; genes with adjusted p-values < 0.05 were considered significant. SVA was performed using smartSVA. GSEA used the fgsea package on pre-ranked gene lists, with FDR-corrected NES reported. LOESS trajectory analysis was adapted from Piehl et al. 2022^69^. Statistical comparisons of physiological measurements (LH, core temperature) and immunofluorescence quantification used one-way ANOVA with Tukey’s post-hoc test or Mann-Whitney U test as appropriate. P < 0.05 or FDR < 0.05 was considered statistically significant.

## Supporting information

Supplementary figures (combined)

## Author contributions

Conceptualization: JCBB, ETJ, SAP, VMN; Methodology: JCBB, ETJ, SAP, LAS, ANF; Investigation: JCBB, ETJ, SAP; Visualization: JCBB, ETJ, SAP; Supervision: DCP, VMN; Writing—original draft: JCBB, VMN; Writing—review and editing: JCBB, ETJ, SAP, HJ, DCP, VMN

## Data and materials availability

Raw RNA-seq data from mouse posterior hypothalamus and preoptic area regions have been deposited to GEO, project #GSE288245. This paper analyzes existing, publicly available GTEx consortium data available on the GTEx Portal (https://www.gtexportal.org/home/). Original code to perform analyses has been deposited at Github (https://github.com/jcbloom/hypo_menopauseNavarro). Any additional information required to reanalyze the data reported in this study is available from the lead contact upon request.

## Supplementary Material

**Supplementary Figure 1**. Cell type deconvolution of the mouse PH and POA RNA-seq dataset using BisqueRNA, using established marker genes from Hajdarovic et al. 2022^20^.

**Supplementary Figure 2**. STRING analysis of TNF-alpha hallmark pathway leading edge genes in the POA between a) 4-months post-OVX vs. intact and b) 14-days post-OVX vs. intact models.

**Supplementary Figure 3**. Volcano plots highlighting significantly differentially expressed genes between labeled conditions and regions. Genes with FDR < 0.05 are labeled in pink. a) PH 4 months post-OVX vs. 14 days post-OVX (35 protein-coding DEGs), b) POA 4 months post-OVX vs. 14 days post-OVX (15 protein-coding DEGs).

**Supplementary Figure 4**. Cell type deconvolution of the human RNA-seq dataset using BisqueRNA. a) Cell type composition using established marker genes from Hajdarovic et al. 2022^20^. b) Presence of cell types expressing key transcription factors from the human hypothalamus reference atlas (Tadross et al. 2025)^30^.

**Supplementary Figure 5**. LOESS expression trajectories of additional postmortem tissue biomarker genes (*PGR*, *GPER1*, *ESR2*) examined in Tickerhoof et al. 2025^35^.

**Supplementary Figure 6**. Spatial transcriptomic visualization of *Akap5* (a) and *Cdkn1a* (b) from the Allen Brain Atlas Brain Knowledge Platform. Expression localization shown with darker coloring indicating regions with higher expression.

**Supplementary Figure 7**. Expression of previously reported aging-associated genes from the literature in the human female hypothalamus. a) Heatmap of aging-associated genes and their expression across age groups. b) Expression of reported brain-associated aging genes across age.

**Supplementary Figure 8**. Overlap of expressed genes between human and mouse hypothalamus tissue; p-value from hypergeometric test.

**Supplementary Table 1.** Gene set enrichment results (mouse).

**Supplementary Table 2.** Differential gene expression output from DESeq2 (mouse).

**Supplementary Table 3.** Normalized gene expression counts (mouse).

**Supplementary Table 4.** GTEx donor IDs used in this study.

**Supplementary Table 5.** Marker genes used to annotate cell types (human and mouse).

**Supplementary Table 6.** LOESS regression values across age per gene.

**Supplementary Table 7.** Gene set enrichment results (human).

**Supplementary Table 8.** Differential gene expression output from DESeq2 (human).

**Supplementary Table 9.** Normalized gene expression counts (human).

**Supplementary Table 10.** Comparison of GSEA results between human, OVX mouse models (this study), and aged mice (Hajdarovic et al. 2022).

**Supplementary Table 11.** *TACR3/Tacr3* correlated expression ranked gene list.

## References

1. Davis SR, Lambrinoudaki I, Lumsden M, et al. Menopause. Nature Reviews Disease Primers. 2015/04/23 2015;1(1):15004. doi:10.1038/nrdp.2015.4

2. Brinton RD, Yao J, Yin F, Mack WJ, Cadenas E. Perimenopause as a neurological transition state. Nature reviews Endocrinology. Jul 2015;11(7):393–405. doi:10.1038/nrendo.2015.82

3. Herbison AE. Physiology of the GnRH neuronal network. In: Knobil JNaE, ed. Physiology of Reproduction. Academic Press; 2006:1415–1482.

4. Rance NE, Uswandi SV. Gonadotropin-releasing hormone gene expression is increased in the medial basal hypothalamus of postmenopausal women. J Clin Endocrinol Metab. Oct 1996;81(10):3540–6. doi:10.1210/jcem.81.10.8855798

5. Ameratunga D, Goldin J, Hickey M. Sleep disturbance in menopause. Intern Med J. Jul 2012;42(7):742–7. doi:10.1111/j.1445-5994.2012.02723.x

6. Augoulea A, Moros M, Lykeridou A, Kaparos G, Lyberi R, Panoulis K. Psychosomatic and vasomotor symptom changes during transition to menopause. Prz Menopauzalny. Jun 2019;18(2):110–115. doi:10.5114/pm.2019.86835

7. Coborn J, de Wit A, Crawford S, et al. Disruption of Sleep Continuity During the Perimenopause: Associations with Female Reproductive Hormone Profiles. J Clin Endocrinol Metab. Sep 28 2022;107(10):e4144–e4153. doi:10.1210/clinem/dgac447

8. Cohn AY, Grant LK, Nathan MD, et al. Effects of Sleep Fragmentation and Estradiol Decline on Cortisol in a Human Experimental Model of Menopause. J Clin Endocrinol Metab. Oct 18 2023;108(11):e1347–e1357. doi:10.1210/clinem/dgad285

9. Greendale GA, Karlamangla AS, Maki PM. The Menopause Transition and Cognition. JAMA. Apr 21 2020;323(15):1495–1496. doi:10.1001/jama.2020.1757

10. Kravitz HM, Joffe H. Sleep during the perimenopause: a SWAN story. Obstet Gynecol Clin North Am. Sep 2011;38(3):567–86. doi:10.1016/j.ogc.2011.06.002

11. Landis CA, Moe KE. Sleep and menopause. Nurs Clin North Am. Mar 2004;39(1):97–115. doi:10.1016/j.cnur.2003.11.006

12. Reed SD, Newton KM, LaCroix AZ, Grothaus LC, Ehrlich K. Night sweats, sleep disturbance, and depression associated with diminished libido in late menopausal transition and early postmenopause: baseline data from the Herbal Alternatives for Menopause Trial (HALT). Am J Obstet Gynecol. Jun 2007;196(6):593 e1-7; discussion 593 e7. doi:10.1016/j.ajog.2007.03.008

13. Freeman EW, Sammel MD, Lin H, Liu Z, Gracia CR. Duration of menopausal hot flushes and associated risk factors. Obstet Gynecol. May 2011;117(5):1095–1104. doi:10.1097/AOG.0b013e318214f0de

14. Koysombat K, McGown P, Nyunt S, Abbara A, Dhillo WS. New advances in menopause symptom management. Best Pract Res Clin Endocrinol Metab. Jan 2024;38(1):101774. doi:10.1016/j.beem.2023.101774

15. Prague JK, Roberts RE, Comninos AN, et al. Neurokinin 3 receptor antagonism as a novel treatment for menopausal hot flushes: a phase 2, randomised, double-blind, placebo-controlled trial. . Lancet. 2017;389:1809–1820.

16. Christ JP, Navarro VM, Reed SD. Nonhormonal Therapies for Menopausal Vasomotor Symptoms. JAMA. Oct 3 2023;330(13):1278–1279. doi:10.1001/jama.2023.15965

17. Koebele SV, Bimonte-Nelson HA. Modeling menopause: The utility of rodents in translational behavioral endocrinology research. Maturitas. May 2016;87:5–17. doi:10.1016/j.maturitas.2016.01.015

18. Avis NE, Crawford SL, Greendale G, et al. Duration of menopausal vasomotor symptoms over the menopause transition. JAMA Intern Med. Apr 2015;175(4):531–9. doi:10.1001/jamainternmed.2014.8063

19. Soares AG, Kilpi F, Fraser A, et al. Longitudinal changes in reproductive hormones through the menopause transition in the Avon Longitudinal Study of Parents and Children (ALSPAC). Sci Rep. Dec 4 2020;10(1):21258. doi:10.1038/s41598-020-77871-9

20. Hajdarovic KH, Yu D, Hassell LA, et al. Single-cell analysis of the aging female mouse hypothalamus. Nat Aging. Jul 2022;2(7):662–678. doi:10.1038/s43587-022-00246-4

21. Korotkevich G, Sukhov V, Budin N, Shpak B, Artyomov MN, Sergushichev A. Fast gene set enrichment analysis. bioRxiv. 2021:060012. doi:10.1101/060012

22. Szklarczyk D, Gable AL, Nastou KC, et al. The STRING database in 2021: customizable protein-protein networks, and functional characterization of user-uploaded gene/measurement sets. Nucleic Acids Res. Jan 8 2021;49(D1):D605–D612. doi:10.1093/nar/gkaa1074

23. Blomqvist A, Engblom D. Neural Mechanisms of Inflammation-Induced Fever. Neuroscientist. Aug 2018;24(4):381–399. doi:10.1177/1073858418760481

24. Shionoya K, Eskilsson A, Blomqvist A. Prostaglandin production selectively in brain endothelial cells is both necessary and sufficient for eliciting fever. Proceedings of the National Academy of Sciences of the United States of America. Oct 25 2022;119(43):e2122562119. doi:10.1073/pnas.2122562119

25. Love MI, Huber W, Anders S. Moderated estimation of fold change and dispersion for RNA-seq data with DESeq2. Genome Biol. 2014;15(12):550. doi:10.1186/s13059-014-0550-8

26. Smith JT, Cunningham MJ, Rissman EF, Clifton DK, Steiner RA. Regulation of Kiss1 gene expression in the brain of the female mouse. Research Support, N.I.H., Extramural Research Support, U.S. Gov’t, Non-P.H.S. Research Support, U.S. Gov’t, P.H.S. Endocrinology. Sep 2005;146(9):3686–92. doi:10.1210/en.2005-0488

27. Goodman RL, Herbison AE, Lehman MN, Navarro VM. Neuroendocrine control of gonadotropin-releasing hormone: Pulsatile and surge modes of secretion. Journal of neuroendocrinology. May 2022;34(5):e13094. doi:10.1111/jne.13094

28. Consortium GT. The Genotype-Tissue Expression (GTEx) project. Nat Genet. Jun 2013;45(6):580–5. doi:10.1038/ng.2653

29. Chen J, Behnam E, Huang J, et al. Fast and robust adjustment of cell mixtures in epigenome-wide association studies with SmartSVA. BMC Genomics. May 26 2017;18(1):413. doi:10.1186/s12864-017-3808-1

30. Tadross JA, Steuernagel L, Dowsett GKC, et al. A comprehensive spatio-cellular map of the human hypothalamus. Nature. Mar 2025;639(8055):708–716. doi:10.1038/s41586-024-08504-8

31. El Khoudary SR, Greendale G, Crawford SL, et al. The menopause transition and women’s health at midlife: a progress report from the Study of Women’s Health Across the Nation (SWAN). Menopause. Oct 2019;26(10):1213–1227. doi:10.1097/gme.0000000000001424

32. Rance NE. Menopause and the human hypothalamus: evidence for the role of kisspeptin/neurokinin B neurons in the regulation of estrogen negative feedback. Peptides. Jan 2009;30(1):111–22. doi:10.1016/j.peptides.2008.05.016

33. Rometo AM, Krajewski SJ, Voytko ML, Rance NE. Hypertrophy and increased kisspeptin gene expression in the hypothalamic infundibular nucleus of postmenopausal women and ovariectomized monkeys. J Clin Endocrinol Metab. Jul 2007;92(7):2744–50. doi:10.1210/jc.2007-0553

34. Butler T, Glodzik L, Wang XH, et al. Positron Emission Tomography reveals age-associated hypothalamic microglial activation in women. Sci Rep. Aug 3 2022;12(1):13351. doi:10.1038/s41598-022-17315-8

35. Tickerhoof M, Cham H, Ouldibbat L, et al. Postmortem tissue biomarkers of menopausal transition. Mol Psychiatry. Feb 2026;31(2):819–834. doi:10.1038/s41380-025-03177-9

36. Tacutu R, Thornton D, Johnson E, et al. Human Ageing Genomic Resources: new and updated databases. Nucleic Acids Res. Jan 4 2018;46(D1):D1083–d1090. doi:10.1093/nar/gkx1042

37. Bateman RJ, Xiong C, Benzinger TL, et al. Clinical and biomarker changes in dominantly inherited Alzheimer’s disease. The New England journal of medicine. Aug 30 2012;367(9):795–804. doi:10.1056/NEJMoa1202753

38. Diaz-Asper CM, Weinberger DR, Goldberg TE. Catechol-O-methyltransferase polymorphisms and some implications for cognitive therapeutics. NeuroRx. Jan 2006;3(1):97–105. doi:10.1016/j.nurx.2005.12.010

39. Herskovits AZ, Guarente L. SIRT1 in neurodevelopment and brain senescence. Neuron. Feb 5 2014;81(3):471–83. doi:10.1016/j.neuron.2014.01.028

40. Kim J, Basak JM, Holtzman DM. The role of apolipoprotein E in Alzheimer’s disease. Neuron. Aug 13 2009;63(3):287–303. doi:10.1016/j.neuron.2009.06.026

41. O’Day DH. Calmodulin Binding Proteins and Alzheimer’s Disease: Biomarkers, Regulatory Enzymes and Receptors That Are Regulated by Calmodulin. International journal of molecular sciences. Oct 5 2020;21(19)doi:10.3390/ijms21197344

42. Pfeilschifter J, Koditz R, Pfohl M, Schatz H. Changes in proinflammatory cytokine activity after menopause. Endocr Rev. Feb 2002;23(1):90–119. doi:10.1210/edrv.23.1.0456

43. Crandall CJ, Manson JE, Hohensee C, et al. Association of genetic variation in the tachykinin receptor 3 locus with hot flashes and night sweats in the Women’s Health Initiative Study. Menopause. Mar 2017;24(3):252–261. doi:10.1097/gme.0000000000000763

44. Pascual O, Ben Achour S, Rostaing P, Triller A, Bessis A. Microglia activation triggers astrocyte-mediated modulation of excitatory neurotransmission. Proceedings of the National Academy of Sciences of the United States of America. Jan 24 2012;109(4):E197–205. doi:10.1073/pnas.1111098109

45. Aten S, Lynch N, Saper CB, Machado NLS. A brain-body perspective on thermoregulatory adaptation. Curr Biol. Oct 20 2025;35(20):R1016–R1028. doi:10.1016/j.cub.2025.09.023

46. Nakamura Y, Yahiro T, Fukushima A, Kataoka N, Hioki H, Nakamura K. Prostaglandin EP3 receptor-expressing preoptic neurons bidirectionally control body temperature via tonic GABAergic signaling. Sci Adv. Dec 23 2022;8(51):eadd5463. doi:10.1126/sciadv.add5463

47. Arruda AP, Milanski M, Velloso LA. Hypothalamic inflammation and thermogenesis: the brown adipose tissue connection. J Bioenerg Biomembr. Feb 2011;43(1):53–8. doi:10.1007/s10863-011-9325-z

48. Zhang G, Li J, Purkayastha S, et al. Hypothalamic programming of systemic ageing involving IKK-beta, NF-kappaB and GnRH. Nature. May 9 2013;497(7448):211–6. doi:10.1038/nature12143

49. He Z, Xie L, Liu J, Wei X, Zhang W, Mei Z. Novel insight into the role of A-kinase anchoring proteins (AKAPs) in ischemic stroke and therapeutic potentials. Biomed Pharmacother. Jun 2024;175:116715. doi:10.1016/j.biopha.2024.116715

50. Koster KP, Fyke Z, Nguyen TTA, et al. Akap5 links synaptic dysfunction to neuroinflammatory signaling in a mouse model of infantile neuronal ceroid lipofuscinosis. Front Synaptic Neurosci. 2024;16:1384625. doi:10.3389/fnsyn.2024.1384625

51. Sanderson JL, Dell’Acqua ML. AKAP signaling complexes in regulation of excitatory synaptic plasticity. Neuroscientist. Jun 2011;17(3):321–36. doi:10.1177/1073858410384740

52. Weisenhaus M, Allen ML, Yang L, et al. Mutations in AKAP5 disrupt dendritic signaling complexes and lead to electrophysiological and behavioral phenotypes in mice. PloS one. Apr 23 2010;5(4):e10325. doi:10.1371/journal.pone.0010325

53. Engeland K. Cell cycle regulation: p53-p21-RB signaling. Cell Death Differ. May 2022;29(5):946–960. doi:10.1038/s41418-022-00988-z

54. Chinta SJ, Woods G, Rane A, Demaria M, Campisi J, Andersen JK. Cellular senescence and the aging brain. Exp Gerontol. Aug 2015;68:3–7. doi:10.1016/j.exger.2014.09.018

55. Wagner KD, Wagner N. The Senescence Markers p16INK4A, p14ARF/p19ARF, and p21 in Organ Development and Homeostasis. Cells. Jun 19 2022;11(12)doi:10.3390/cells11121966

56. Wang L, Lankhorst L, Bernards R. Exploiting senescence for the treatment of cancer. Nat Rev Cancer. Jun 2022;22(6):340–355. doi:10.1038/s41568-022-00450-9

57. Hall JE. Neuroendocrine physiology of the early and late menopause. Endocrinol Metab Clin North Am. Dec 2004;33(4):637–59. doi:10.1016/j.ecl.2004.08.002

58. Hall JE, Lavoie HB, Marsh EE, Martin KA. Decrease in gonadotropin-releasing hormone (GnRH) pulse frequency with aging in postmenopausal women. J Clin Endocrinol Metab. May 2000;85(5):1794–800. doi:10.1210/jcem.85.5.6612

59. Lohff JC, Christian PJ, Marion SL, Hoyer PB. Effect of duration of dosing on onset of ovarian failure in a chemical-induced mouse model of perimenopause. Menopause. May-Jun 2006;13(3):482–8. doi:10.1097/01.gme.0000191883.59799.2e

60. Mayer LP, Devine PJ, Dyer CA, Hoyer PB. The follicle-deplete mouse ovary produces androgen. Biology of reproduction. Jul 2004;71(1):130–8. doi:10.1095/biolreprod.103.016113

61. Van Kempen TA, Milner TA, Waters EM. Accelerated ovarian failure: a novel, chemically induced animal model of menopause. Brain research. Mar 16 2011;1379:176–87. doi:10.1016/j.brainres.2010.12.064

62. Cao LB, Leung CK, Law PW, et al. Systemic changes in a mouse model of VCD-induced premature ovarian failure. Life Sci. Dec 1 2020;262:118543. doi:10.1016/j.lfs.2020.118543

63. Cao LB, Liu HB, Lu G, et al. Hormone-Like Effects of 4-Vinylcyclohexene Diepoxide on Follicular Development. Front Cell Dev Biol. 2020;8:587. doi:10.3389/fcell.2020.00587

64. Kreisman MJ, McCosh RB, Tian K, Song CI, Breen KM. Estradiol Enables Chronic Corticosterone to Inhibit Pulsatile Luteinizing Hormone Secretion and Suppress Kiss1 Neuronal Activation in Female Mice. Neuroendocrinology. 2020;110(6):501–516. doi:10.1159/000502978

65. Wang Q, Ding SL, Li Y, et al. The Allen Mouse Brain Common Coordinate Framework: A 3D Reference Atlas. Cell. May 14 2020;181(4):936–953 e20. doi:10.1016/j.cell.2020.04.007

66. Bray NL, Pimentel H, Melsted P, Pachter L. Near-optimal probabilistic RNA-seq quantification. Nat Biotechnol. May 2016;34(5):525–7. doi:10.1038/nbt.3519

67. Soneson C, Love MI, Robinson MD. Differential analyses for RNA-seq: transcript-level estimates improve gene-level inferences. F1000Research. 2015;4:1521. doi:10.12688/f1000research.7563.2

68. Frankish A, Diekhans M, Ferreira AM, et al. GENCODE reference annotation for the human and mouse genomes. Nucleic Acids Res. Jan 8 2019;47(D1):D766–D773. doi:10.1093/nar/gky955

69. Piehl N, van Olst L, Ramakrishnan A, et al. Cerebrospinal fluid immune dysregulation during healthy brain aging and cognitive impairment. Cell. Dec 22 2022;185(26):5028–5039 e13. doi:10.1016/j.cell.2022.11.019

70. Jew B, Alvarez M, Rahmani E, et al. Accurate estimation of cell composition in bulk expression through robust integration of single-cell information. Nat Commun. Apr 24 2020;11(1):1971. doi:10.1038/s41467-020-15816-6

