## Supplementary figures (combined) for "Progressive hypothalamic neuroinflammation after ovariectomy in mice parallels age-related transcriptomic changes in the female human hypothalamus"

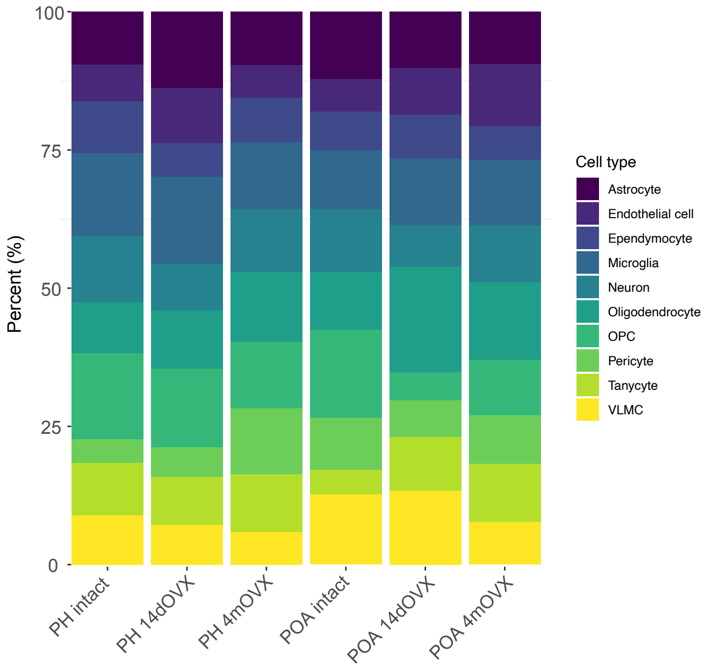


Supplementary Figure 1

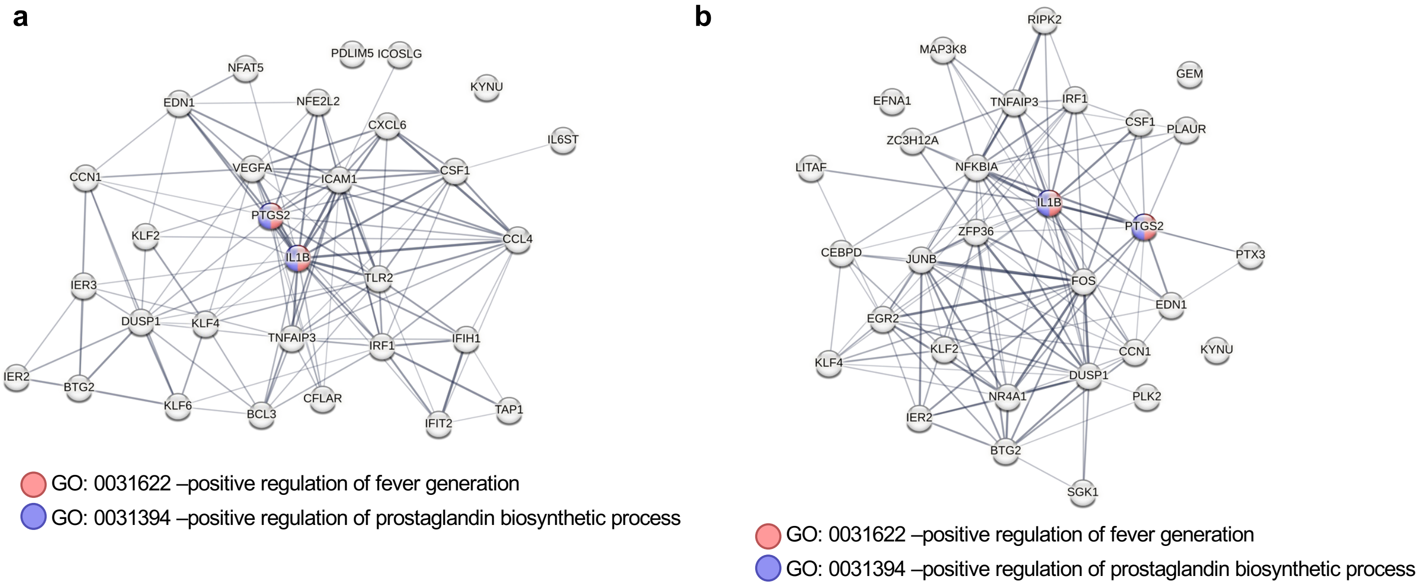


Supplementary Figure 2

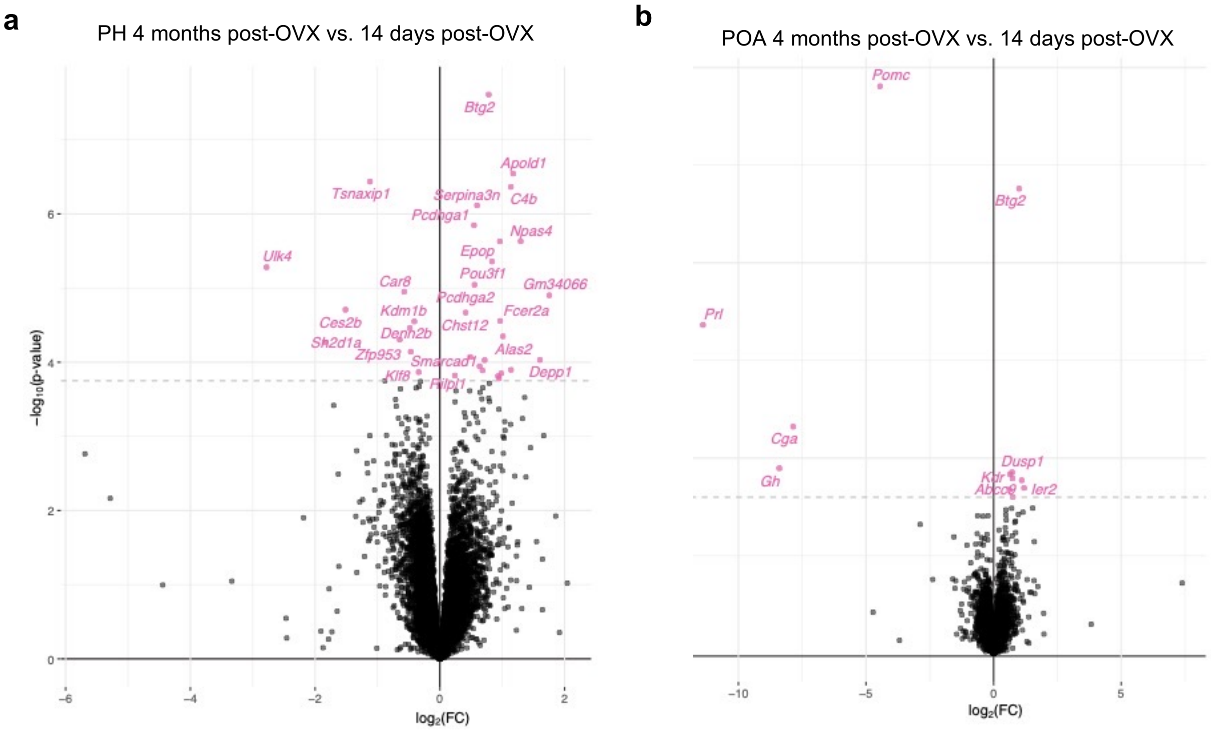


Supplementary Figure 3

a

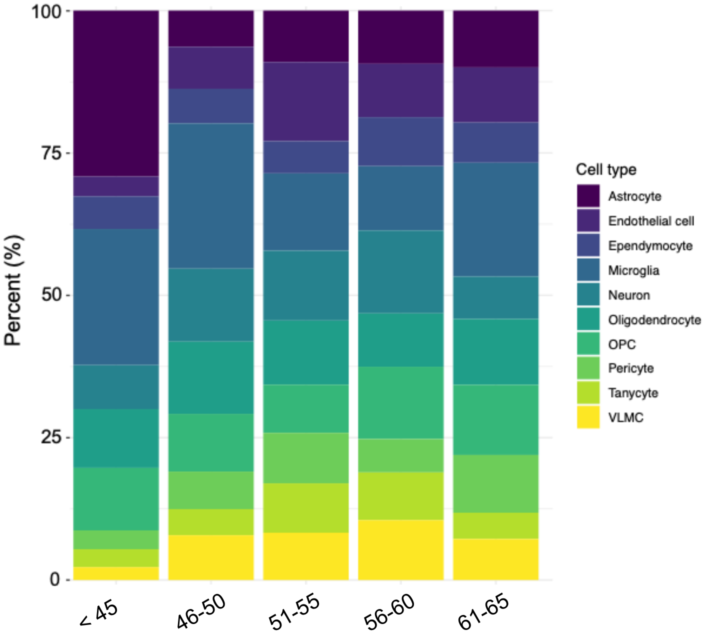


b

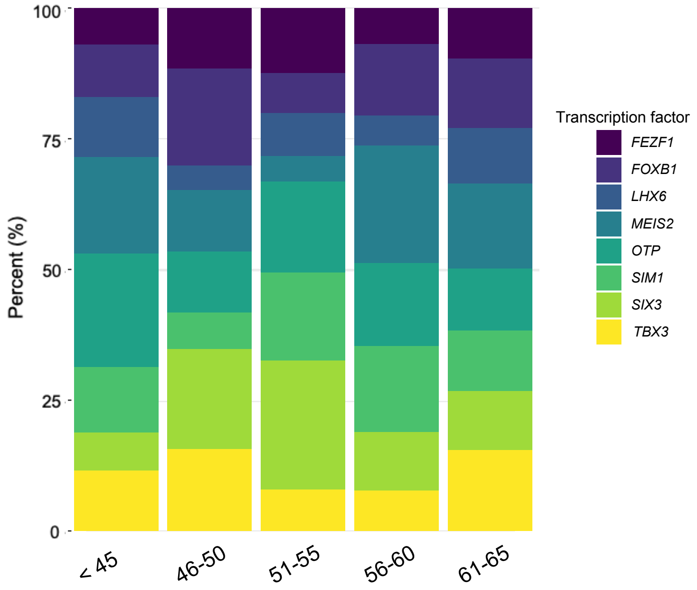


Supplementary Figure 4

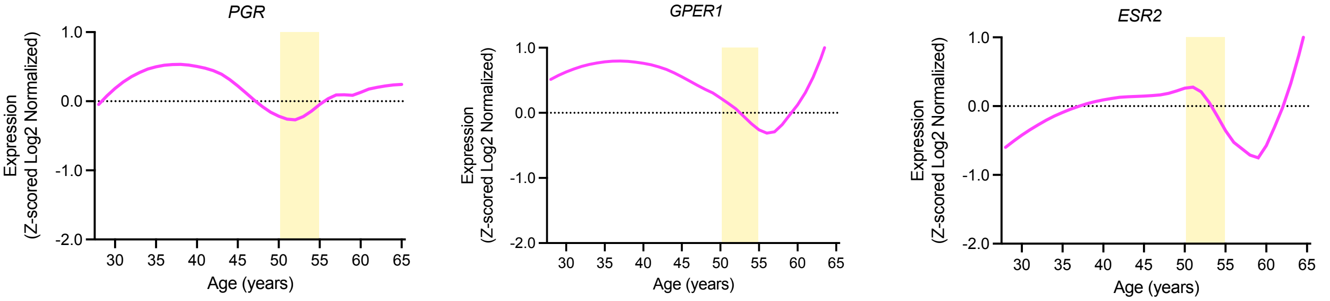


Supplementary Figure 5

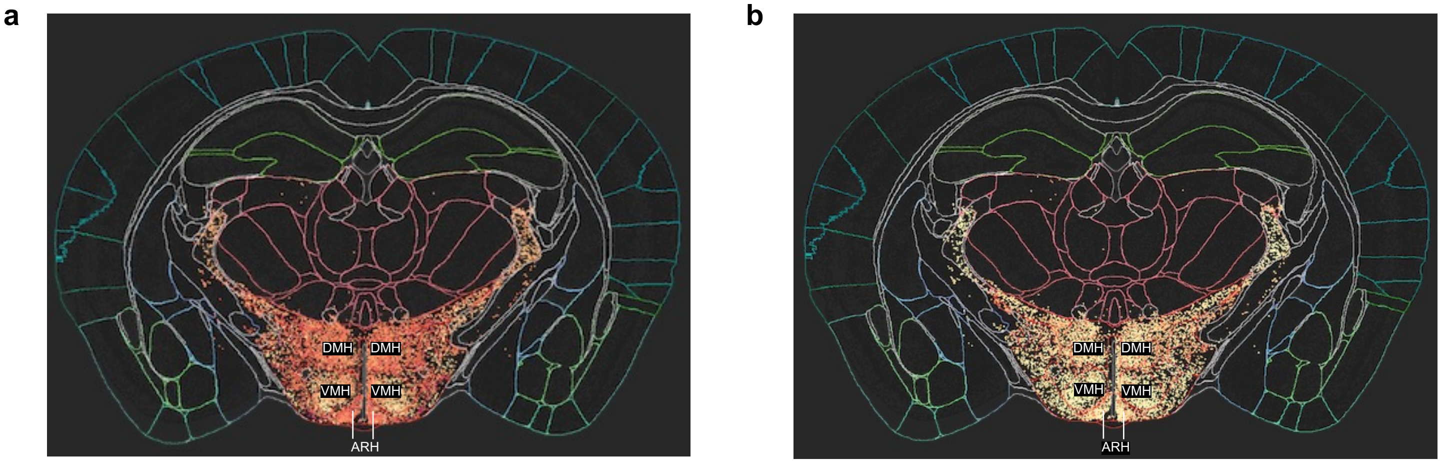


Supplementary Figure 6

a

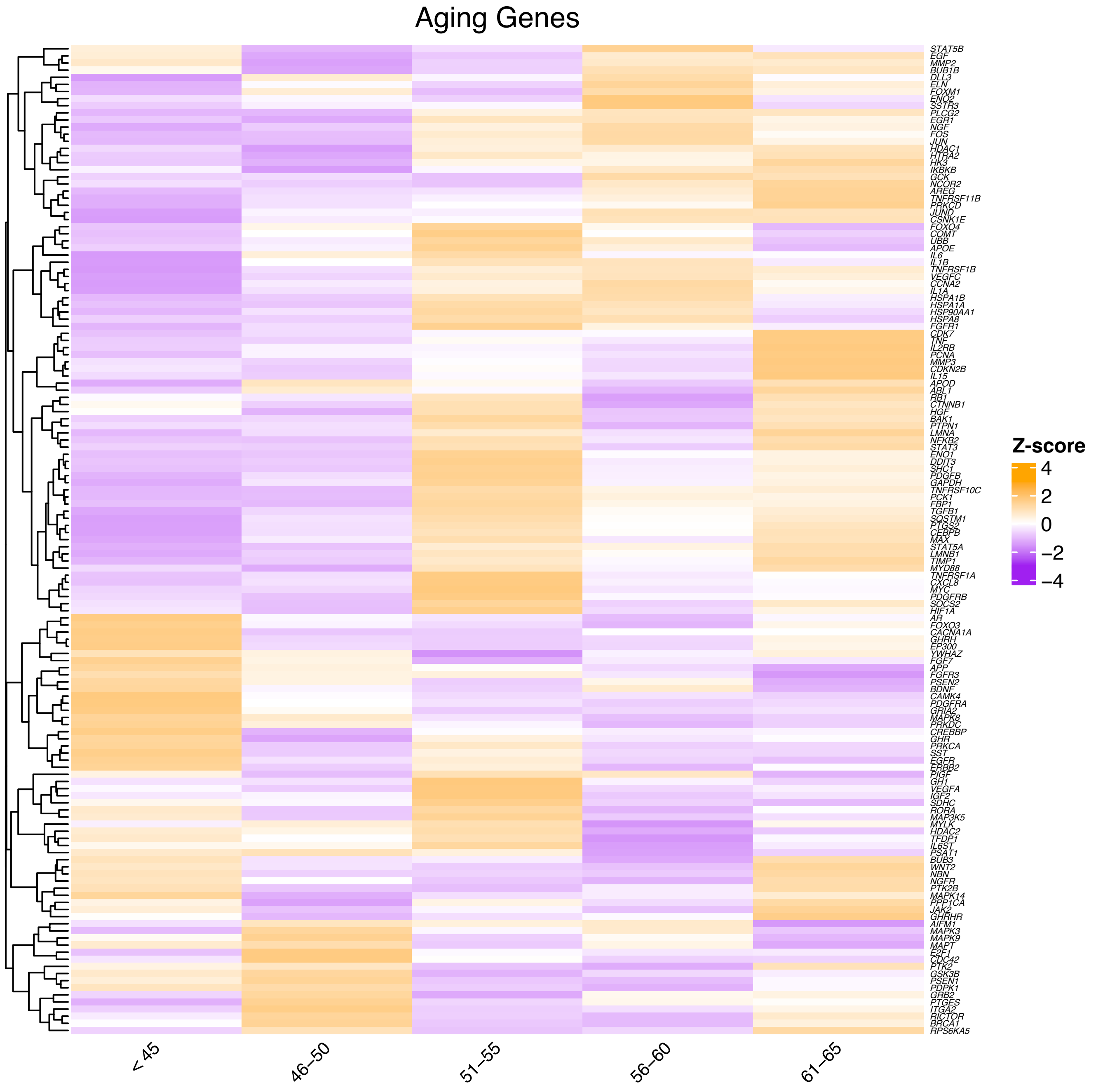


b

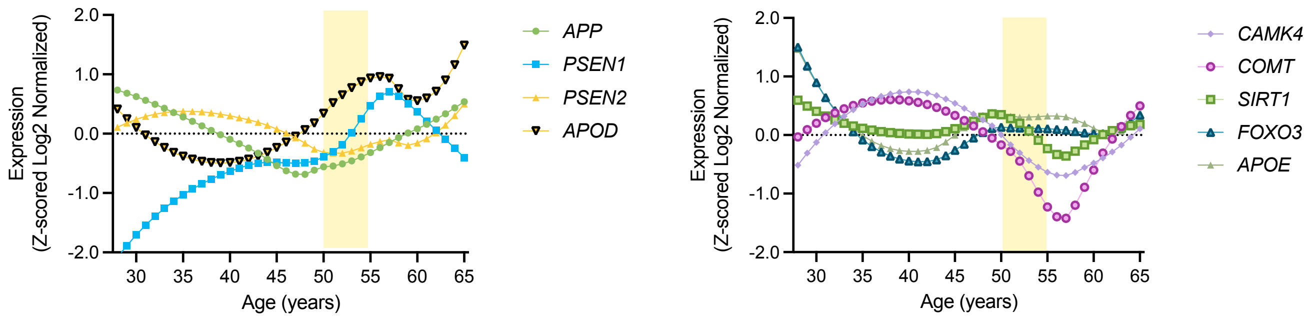


Supplementary Figure 7

a

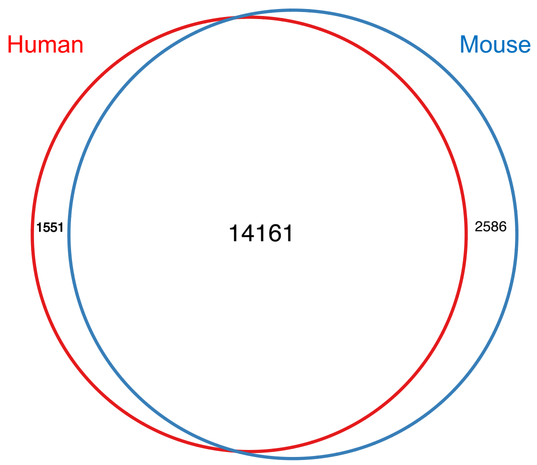


Supplementary Figure 8
